# Risk factors associated with canine overweightness and obesity in an owner-reported survey

**DOI:** 10.1101/2020.01.06.896399

**Authors:** LeeAnn M. Perry, Justin Shmalberg, Jirayu Tanprasertsuk, Dan Massey, Ryan W. Honaker, Aashish R. Jha

## Abstract

**Background:** Overweightness and obesity in dogs are associated with negative health outcomes. A better understanding of risk factors associated with canine weight is fundamental to identifying preventative interventions and treatments. In this cross-sectional study, we used a direct to consumer approach to collect body condition scores (BCS), as well as demographic, diet, and lifestyle data on 4,446 dogs. BCS was assessed by owners using a 9-point system and categorized as ideal (BCS 4-5), overweight (BCS 6), and obese (BCS 7+). Following univariate analyses, a stepwise procedure was used to select variables which were included in multivariate logistic regression models. One model was created to compare ideal to all overweight and obese dogs, and another was created to compare ideal to obese dogs only. We then used Elastic Net selection and XGBoost variable importance measures to validate these results.

**Results:** Overall, 1,480 (33%) of dogs were reported to be overweight or obese, of which 356 (8% total) of dogs were reported to be obese. Seven factors were significantly associated with both overweightness/obesity and obesity alone in all three analyses (stepwise, Elastic Net, and XGBoost): diet composition, probiotic supplementation, treat quantity, exercise, age, food motivation level, and pet appetite. Neutering was also associated with overweightness/obesity in all analyses.

**Conclusions:** This study recapitulated established risk factors associated with BCS (age, exercise, neutering). Moreover, we elucidated associations between previously examined risk factors and BCS (diet composition, treat consumption, and temperament) and identified a novel factor (probiotic supplementation). Specifically, relative to dogs on fresh food diets, BCS was higher in dogs eating dry food both alone and in combination with other foods. Furthermore, dogs receiving probiotics, but not other forms of supplementation, were more likely to have an ideal BCS. Future studies should corroborate these findings with experimental manipulations.

## Background

Overweightness and obesity are major health concerns in both humans and companion dogs. Weight issues have been associated with myriad negative health conditions as well as decreased life span (1, 2). In the largest study to date, the prevalence of veterinarian-assessed overweightness and obesity of dogs in the United States has been reported to be 34% and 5% respectively (3). Globally, canine overweightness has been determined to range from 6-31% in European countries, 44% in China, 40% in Japan, and 26% in Australia (4–7).

Clinical assessment of overweightness and obesity relies on body condition scores (BCS) based on visual inspection and palpation. A number of studies have identified risk factors associated with increased BCS, including increased age (3-6,8,9), neutering (3,5,6,9,10), and decreased exercise (11, 12). Some studies have also identified an effect of different feeding practices. Home-made diets, table foods, semi-moist foods, and canned foods have all been associated with overweightness and obesity (3, 12). The consumption of treats and snacks has also been identified as a risk factor (8,10,11). Despite these findings, many of these studies have been limited by comparatively smaller sample sizes (8), or have included only a handful of risk factors (6). Thus, investigating factors associated with canine body weight using a larger cohort and a more comprehensive list of risk factors is warranted.

This study addresses these drawbacks by engaging a direct-to-consumer data collection model wherein a large pool of pet owners distributed widely across the United States completed an extensive online health assessment. The volume of data collected not only allowed us to use standard analytic methods, but also permitted the use of machine learning approaches that do not have the same biases or assumptions of traditional statistical methods. Our results recapitulate findings from previous studies on canine overweightness and obesity, provide additional evidence for factors with conflicting results in the literature, and identify novel factors associated with overweightness and obesity.

## Results

Out of a total of 4,446 dogs, 2,967 (67%) were at an ideal weight, defined as an owner-reported body condition score of 4-5 on a previously-validated 9-point system (BCS 4-5; (13)) and 1,480 (33%) were overweight or obese (BCS≥6). Of these dogs with higher BCS, 1,124 (25% of total) were overweight (BCS 6) and 356 (8% of total) were obese (BCS≥7).

### Identification of Risk Factors Associated with Overweightness and Obesity

#### Significant Risk Factors Identified via Univariate Analysis

To identify factors associated with increased BCS, we performed two univariate analyses comparing ideal weight dogs (N=2,966) to overweight/obese dogs (N=1,480) as well as to obese dogs only (N=356). Of the 45 variables selected as outlined in Methods, 22 (49%) were significantly associated with overweightness/obesity in a univariate analysis (p < 0.05, N=4,446) and 18 (40%) were significantly associated with obesity (p < 0.05, N=3,322). The relationships between these variables and BCS are summarized in Table 1. The 18 variables positively associated with both overweightness/obesity and obesity were diet combinations containing dry food, increased treat quantity, lack of probiotic consumption, increased age, decreased exercise per week, neutering, increased pet appetite, increased food motivation, lower overall mood, decreased conspecific interaction, increased tail chasing, decreased prey drive, presence of other dogs in the household, rural home environment, conventional-only medicine type, household tobacco use, food intolerances, and rescue or other acquisition method. Four additional variables positively associated with BCS in the overweightness/obesity analysis were the use of dental chews, sharing or cooking food, increased overall nervousness, and decreased dental visit frequency.

**Table 1:** Results of Univariate Analyses.

| Table 1: Results of Univariate Analyses |  |  |  |  |  |  |  |
| --- | --- | --- | --- | --- | --- | --- | --- |
|  |  | Overweight and Obese (N=1480) vs Ideal (N=2966) |  |  | Obese (N=356) vs Ideal (N=2966) |  |  |
|  |  | N (% of 4,446) or mean (SD) | OR [95% CI] | p | N (% of 3,322) or mean (SD) | OR [95% CI] | p |
| Diet | Fresh Only | 1,001 (22%) | Reference |  | 761 (23%) | Reference |  |
|  | Kibble Only | 938 (21%) | 1.45 [1.21-1.76] | <0.0001*** | 684 (21%) | 2.11 [1.52-2.96] | <0.0001*** |
|  | Kibble & Fresh | 376 (8.5%) | 1.31 [1.02-1.68] | 0.03* | 263 (7.9%) | 1.08 [0.64-1.76] | >0.05 |
|  | Kibble & Canned | 349 (7.9%) | 1.85 [1.44-2.38] | <0.0001*** | 238 (7.2%) | 2.56 [1.68-3.87] | <0.0001*** |
|  | Canned Only | 182 (4.1%) | 1.31 [0.94-1.83] | >0.05 | 134 (4.0%) | 1.75 [0.97-3.01] | >0.05 |
|  | Raw Only | 136 (3.1%) | 0.52 [0.32-0.81] | 0.005** | 116 (3.5%) | 0.51 [0.17-1.17] | >0.05 |
|  | Dried Only | 109 (2.5%) | 0.84 [0.53-1.30] | >0.05 | 86 (2.6%) | 0.85 [0.32-1.87] | >0.05 |
|  | Other/Unknown | 1,355 (30%) | 0.98 [0.82-1.18] | >0.05 | 1,040 (31%) | 1.06 [0.75-1.49] | >0.05 |
| Treats | None | 325 (7.3%) | Reference |  | 262 (7.9%) | Reference |  |
|  | <10 percent | 3,535 (80%) | 1.13 [0.88-1.46] | >0.05 | 2,655 (80%) | 0.72 [0.49-1.09] | >0.05 |
|  | >10 percent | 581 (13%) | 2.12 [1.59-2.84] | <0.0001*** | 402 (12%) | 2.15 [1.40-3.39] | 0.0007*** |
| Probiotics | No | 3,602 (81%) | Reference |  | 2,658 (80%) | Reference |  |
|  | Yes | 844 (19%) | 0.65 [0.55-0.77] | <0.0001*** | 664 (20%) | 0.46 [0.32-0.64] | <0.0001*** |
| Age | Years | 6.73 (4.09) | 1.09 [1.07-1.11] | <0.0001*** | 6.44 (4.16) | 1.10 [1.07-1.13] | <0.0001*** |
| Exercise per week | 0-4 hours | 1,827 (41%) | Reference |  | 1,296 (39%) | Reference |  |
|  | 4-7 hours | 1,549 (35%) | 0.57 [0.50-0.66] | <0.0001*** | 1,160 (35%) | 0.28 [0.21-0.37] | <0.0001*** |
|  | 7-14 hours | 804 (18%) | 0.45 [0.37-0.54] | <0.0001*** | 639 (19%) | 0.25 [0.17-0.35] | <0.0001*** |
|  | 14+ hours | 257 (5.8%) | 0.29 [0.20-0.40] | <0.0001*** | 221 (6.7%) | 0.19 [0.09-0.35] | <0.0001*** |
| Neutered | No | 524 (12%) | Reference |  | 449 (14%) | Reference |  |
|  | Yes | 3,904 (88%) | 2.39 [1.90-3.02] | <0.0001*** | 2,864 (86%) | 2.54 [1.67-4.08] | <0.0001*** |
| Pet Appetite | 1-5 scale | 1.90 (0.46) | 2.16 [1.86-2.51] | <0.0001*** | 1.88 (0.46) | 4.30 [3.28-5.67] | <0.0001*** |
| Food Motivation | 1-5 scale | 4.00 (1.26) | 1.23 [1.16-1.30] | <0.0001*** | 3.95 (1.28) | 1.40 [1.26-1.56] | <0.0001*** |
| Mood | 1-5 scale | 1.91 (0.93) | 1.30 [1.21-1.38] | <0.0001*** | 1.88 (0.93) | 1.50 [1.34-1.67] | <0.0001*** |
| Conspecific Interaction | 1-4 scale | 2.67 (0.94) | 0.81 [0.75-0.86] | <0.0001*** | 2.70 (0.94) | 0.71 [0.63-0.81] | <0.0001*** |
| Tail Chasing | 1-5 scale | 4.74 (0.74) | 1.24 [1.13-1.36] | <0.0001*** | 4.73 (0.76) | 1.54 [1.25-1.97] | 0.0001*** |
| Prey Drive | 1-5 scale | 2.67 (1.29) | 0.91 [0.87-0.96] | 0.0003*** | 2.69 (1.30) | 0.83 [0.76-0.91] | <0.0001*** |
| Other Dogs | No | 2,361 (53%) | Reference |  | 1,775 (53%) | Reference |  |
|  | Yes | 2,085 (47%) | 1.26 [1.11-1.43] | 0.0003*** | 1,547 (47%) | 1.81 [1.45-2.27] | <0.0001*** |
| Dental Chews | No | 2,786 (63%) | Reference |  | 2,120 (64%) | Reference |  |
|  | Yes | 1,660 (37%) | 1.24 [1.09-1.41] | 0.001** | 1,202 (36%) | 1.23 [0.98-1.54] | >0.05 |
| Share Food/Cook | No | 1,768 (40%) | Reference |  | 1,362 (41%) | Reference |  |
|  | Yes | 2,668 (60%) | 1.24 [1.09-1.41] | 0.001** | 1,953 (59%) | 1.19 [0.95-1.50] | >0.05 |
| Home Environment | Urban | 1,048 (24%) | Reference |  | 813 (24%) | Reference |  |
|  | Suburban | 2,736 (62%) | 1.22 [1.05-1.43] | 0.009** | 2,017 (61%) | 1.16 [0.88-1.53] | >0.05 |
|  | Rural | 648 (15%) | 1.36 [1.11-1.68] | 0.003** | 480 (14%) | 1.62 [1.14-2.30] | 0.007** |
| Medicine Type | Conventional Only | 1,621 (36%) | Reference |  | 1,202 (36%) | Reference |  |
|  | Mixed | 2,602 (59%) | 0.84 [0.74-0.96] | 0.01* | 1,949 (59%) | 0.65 [0.52-0.82] | 0.0002*** |
|  | Holistic Only | 176 (4.0%) | 0.67 [0.47-0.95] | 0.03* | 138 (4.2%) | 0.51 [0.25-0.94] | 0.04* |
| Nervousness | 1-5 scale | 2.36 (1.21) | 1.06 [1.01-1.12] | 0.02* | 2.34 (1.20) | 1.09 [0.99-1.19] | >0.05 |
| Household Tobacco | No | 4,098 (92%) | Reference |  | 3,070 (92%) | Reference |  |
|  | Yes | 327 (7%) | 1.34 [1.06-1.69] | 0.01* | 236 (7%) | 1.69 [1.16-2.41] | 0.005** |
| Dentist Frequency | 1-3 scale | 1.56 (0.73) | 1.11 [1.02-1.21] | 0.02* | 1.55 (0.73) | 1.12 [0.97-1.30] | >0.05 |
| Food Intolerances | No | 3,646 (82%) | Reference |  | 2,714 (82%) | Reference |  |
|  | Yes | 774 (18%) | 0.83 [0.70-0.98] | 0.03* | 591 (18%) | 0.70 [0.51-0.96] | 0.03* |
| Acquisition Method | Breeder | 1,930 (43%) | Reference |  | 1,470 (44%) | Reference |  |
|  | Rescue | 1,657 (37%) | 1.25 [1.09-1.44] | 0.002** | 1,212 (36%) | 1.38 [1.07-1.78] | 0.0002*** |
|  | Other | 859 (19%) | 1.29 [1.09-1.53] | 0.003** | 640 (19%) | 1.75 [1.31-2.33] | 0.01* |

OR: Odds Ratio; 95% CI: 95% Confidence Interval; Pet Appetite: 1-Poor, 5-Excellent; Food Motivation Level: 1-Not at all, 5-Very; Mood: 1-Excellent, 5-Depressed; Conspecific Interaction: 1-Never, 4-Often; Tail Chasing Frequency: 1-Never, 5-Multiple times per day; Prey Drive: 1-Never, 5-Very Often; Nervousness: 1-Not at all, 5-Very; Dentist Frequency: 1-Never, 3-Three or more times

#### Significant Risk Factors Identified via Stepwise Multivariable Analysis

Since body weight is a complex trait that can be influenced by multiple variables, we performed multivariate analysis to understand how different variables are associated with BCS when they are taken together. We performed these analyses in the overweight/obese and obese groups separately. The results from the two stepwise logistic regression models comparing ideal weight dogs (N=2,725) to overweight/obese dogs (N=1,384) as well as obese dogs (N=327) are presented in Table 2. Due to missing data in the selected variables, 337 dogs from the total dataset were dropped from this logistic regression. Log odds for significant variables are presented in Figure 1. The variables significantly associated with overweightness/obesity were age, exercise per week, food motivation level, overall mood, pet appetite, sharing food, neutering, treat quantity, probiotic supplements, home environment, diet, and dental treatment frequency (p < 0.05, stepwise logistic regression, N=4,109). The variables significantly associated with obesity alone were age, exercise per week, food motivation level, other dogs in the household, overall mood, pet appetite, neutering, tail chasing frequency, treat quantity, probiotic supplements, medicine type, diet, and dental treatment frequency (p < 0.05, stepwise logistic regression, N=3,052). The intersection of variables in the two models were age, exercise per week, food motivation level, overall mood, pet appetite, neutering, treat quantity, probiotic supplements, diet, and dental treatment frequency.

**Figure 1.**
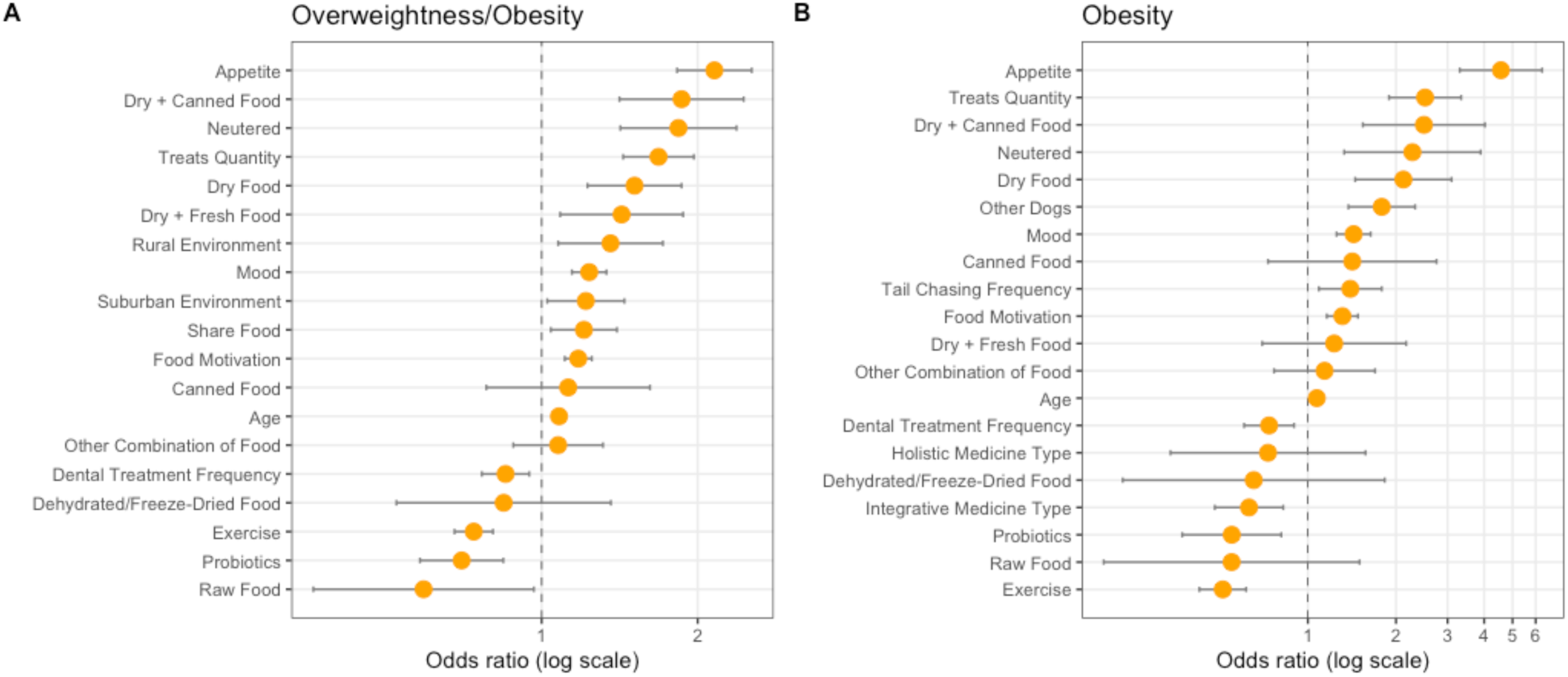
Variable Log Odds in Final Stepwise Models. Odds ratios for each significant variable in the final model for (A) overweightness and obesity and (B) obesity alone shown. Error bars show the 95% confidence interval for each risk factor. Diet risk factors are shown relative to a fresh diet, home environment risk factors are shown relative to an urban environment, and medicine type risk factors are shown relative to conventional medicine only.

**Table 2:**
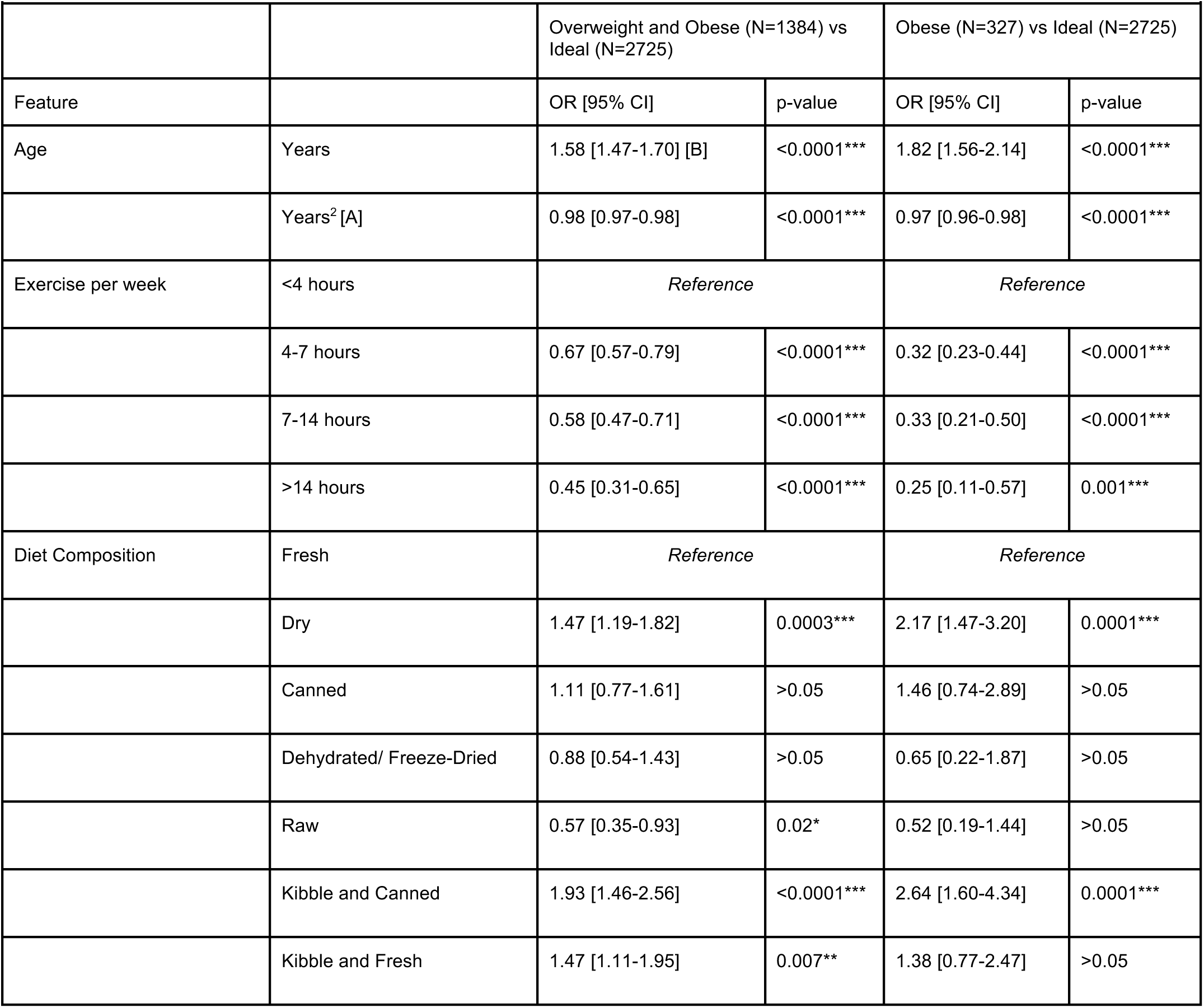
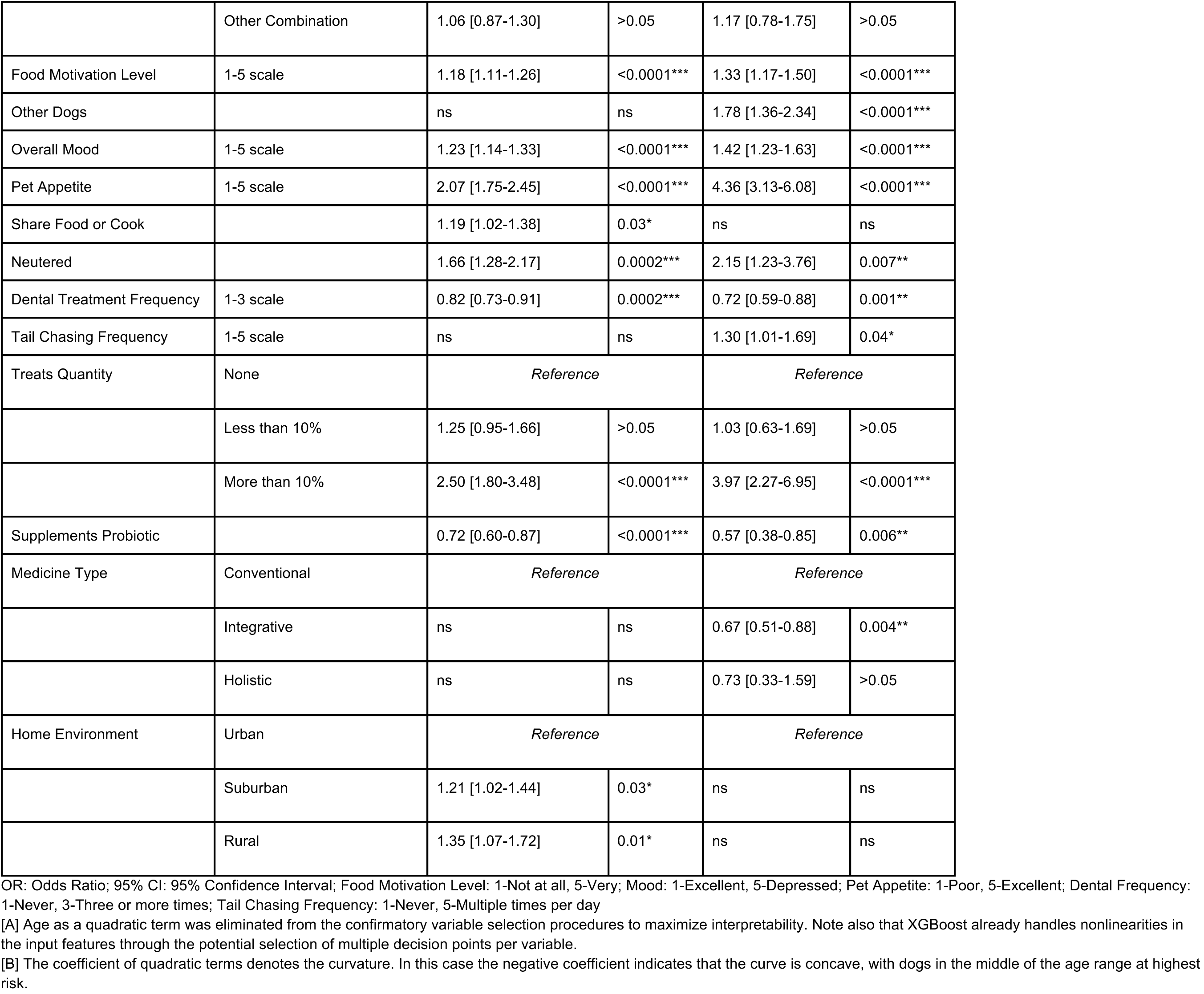
Final Multivariable Logistic Regression Models.

Given the nonlinear relationship between age and body condition reported by others (3,5,6,9), age was entered as both a linear and quadratic term in the logistic regression model. In the context of the model, both the linear and quadratic terms had odds ratios that were statistically significant (p<0.0001). The inclusion of the quadratic term significantly improved both the overweight/obese and obese only logistic regression models, as determined by nested model ANOVA (p<0.0001, likelihood ratio test).

#### Significant Risk Factors Selected via Elastic Net Analysis

Collinearity poses a major issue for stepwise models, since only a subset of a group of collinear variables may be selected, such that the final ensemble of variables may be influenced by noise. Elastic Net is a method that combines the L_1_ and L_2_ penalties used in the Lasso and ridge methods, respectively. It addresses the collinearity issue, as it exhibits a grouping effect (14) wherein coefficients of correlated variables tend to be similar. Thus, through the Elastic Net algorithm we may see if any important variables are being masked by their correlations with others.

For both the overweight/obese and obese only models, the most significant ensemble of risk factors was selected as detailed in Methods. Factors that appeared in both the optimal model for overweightness/obesity and the optimal model for obesity alone were: pet appetite, treat quantity, exercise, probiotic supplementation, diet, mood, food motivation level, and age. The overweightness/obesity model also included neutering, home environment, and sharing food, while the obesity model also included medicine type. With the exception of medicine type, all of these variables are within the subset of those selected by stepwise logistic regression. Variables selected by the Elastic Net are presented in [Supplementary File 1] and [Supplementary File 2]. Comparisons between variables selected by Elastic Net and other methods are presented in Table 3.

**Table 3:** Significant variables selected by each protocol.

|  | Overweightness/Obesity |  |  |  | Obesity |  |  |  |
| --- | --- | --- | --- | --- | --- | --- | --- | --- |
|  | Univariate | Stepwise | Elastic Net | XGBoost | Univariate | Stepwise | Elastic Net | XGBoost |
| Diet | ● | ● | ● | ● | ● | ● | ● | ● |
| Probiotic Supplements | ● | ● | ● | ● | ● | ● | ● | ● |
| Treat Quantity | ● | ● | ● | ● | ● | ● | ● | ● |
| Age | ● | ● | ● | ● | ● | ● | ● | ● |
| Exercise | ● | ● | ● | ● | ● | ● | ● | ● |
| Food Motivation Level | ● | ● | ● | ● | ● | ● | ● | ● |
| Pet Appetite | ● | ● | ● | ● | ● | ● | ● | ● |
| Overall Mood | ● | ● | ● | ○ | ● | ● | ● | ○ |
| Other Dogs in Household | ● | ● | ○ | ○ | ● | ● | ○ | ○ |
| Sharing Food | ● | ● | ● | ○ | ○ | ○ | ○ | ○ |
| Neutered | ● | ● | ● | ● | ● | ● | ○ | ○ |
| Home Environment | ● | ● | ● | ○ | ● | ○ | ○ | ○ |
| Dentist Frequency | ● | ● | ○ | ○ | ○ | ● | ○ | ○ |
| Tail Chasing Frequency | ● | ○ | ○ | ○ | ● | ● | ○ | ○ |
| Medicine Type | ● | ○ | ○ | ○ | ● | ○ | ● | ○ |

● $p < 0.05$ , ○ $p > 0.05$ , Variables were included if they were significant in at least two different models.

#### Significant Risk Factors Selected via XGBoost Analysis

Another issue with the stepwise model is that non-linear effects are not accounted for. We addressed this by using XGBoost, a tree-based machine learning algorithm (15). A tree-based model can establish decision points at multiple different values, and thus the final variable importance in the model encompasses non-linear relationships including interactions between variables. Through XGBoost we may identify which variables, if any, we should examine in terms of higher-order interactions or polynomial models. Factors that appeared in both the optimal model for overweightness/obesity and the optimal model for obesity were: age, pet appetite, exercise, treat quantity, food motivation level, diet, and probiotic consumption. The overweightness/obesity model also included neutering. Each one of these variables was also selected by stepwise logistic regression and Elastic Net. The variable importance plots from the XGBoost models are presented in [Supplementary File 3] and [Supplementary File 4]. Comparisons between variables selected by XGBoost and other methods are presented in Table 3.

#### Healthy Subgroup

We repeated the multivariate logistic regression analysis with only the subgroup of dogs that were reported to have no major health conditions, in order to remove the possibly confounding effects of disease and treatment variables. This subset consisted of dogs that did not have pancreatitis, diabetes, kidney issues, liver disease, heart issues, cancer, or gastrointestinal conditions (N=3,173, 71% of total dataset). Of these dogs, 2,118 (67%) were at an ideal weight (BCS 4-5) and 1,055 (33%) were either overweight/obese (BCS≥6) or obese (BCS≥7); these were further categorized into 792 (25% of total) that were overweight and 263 (8% of total) that were obese. These proportions were not statistically significantly different from the proportions in the overall sample (p < 0.05, χ^2^ test). Thirteen of the 15 variables selected from the total dataset remained significant despite decreased power to detect significant effects. These findings are available in [Supplementary File 5]. Due to missingness in the selected variables, 192 dogs were dropped from the healthy subgroup logistic regression.

### Individual Contributions of Selected Risk Factors

Among the 7 risk factors identified by all of the eight selection methods (univariate, stepwise, Elastic Net, and XGBoost each undertaken for both overweightness/obesity and obesity alone) [Table 3] were diet, age, exercise, probiotic supplementation, and treat quantity. We further examined the individual contributions of these five risk factors to BCS.

#### Diet

We examined the relationship between the most common diet types and both overweightness/obesity (Figure 2) and obesity alone (Figure 3). Since a fresh food only diet was the largest category (besides “Other”, see Methods) we used this as our reference level in the logistic regression.

**Figure 2:**
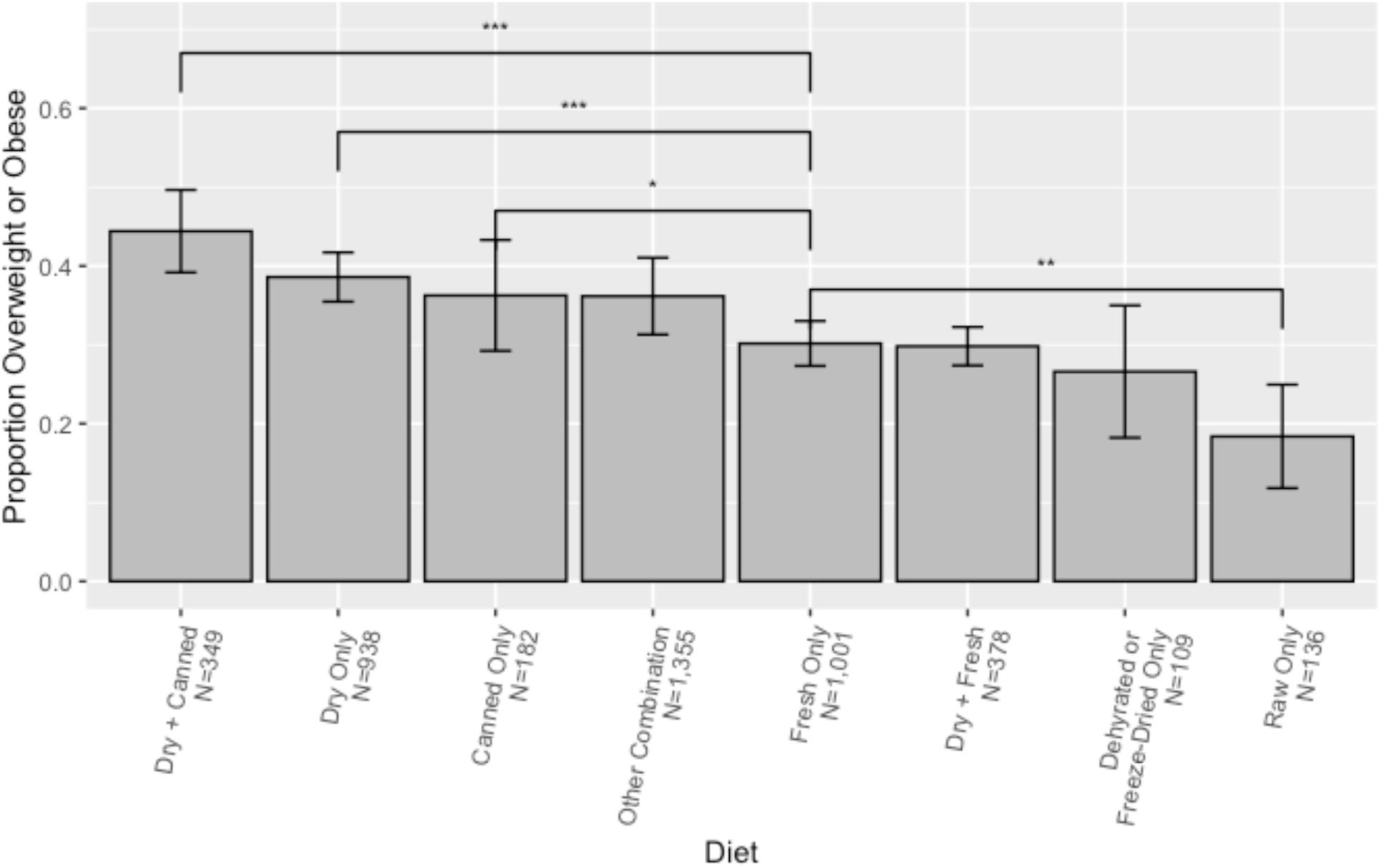
Proportion of Dogs Overweight/Obese Relative to a Fresh Food Diet. Error bars show 95% confidence intervals. *, **, *** respectively represent p<0.05, p<0.01, p<0.001. “Other” includes any combination of reported dietary elements not represented in the most common dietary configurations, for example having three or more regular dietary elements.

**Figure 3:**
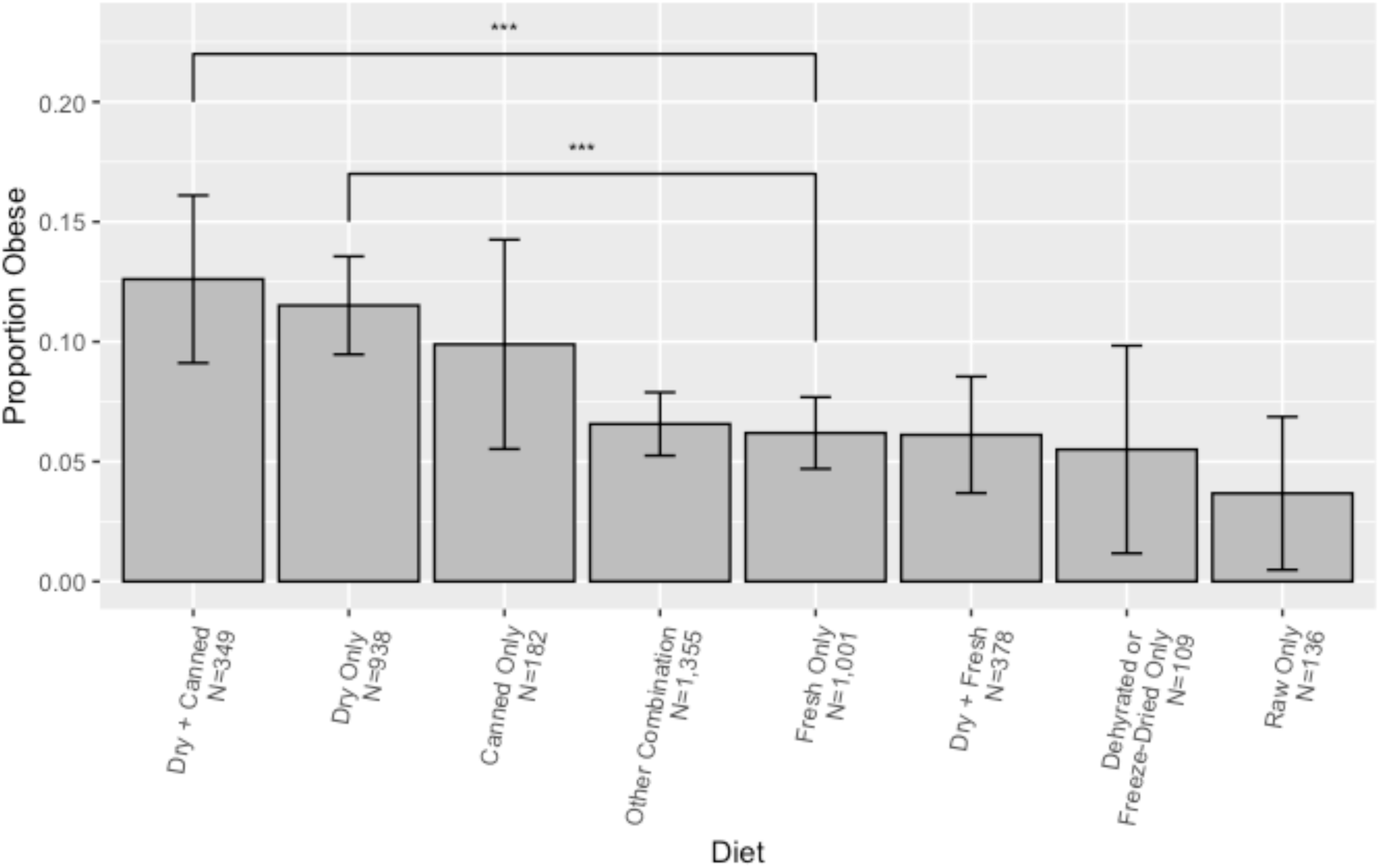
Proportion of Dogs Obese Relative to a Fresh Food Diet. Error bars show 95% confidence intervals. *, **, *** respectively represent p<0.05, p<0.01, p<0.001. “Other” includes any combination of reported dietary elements not represented in the most common dietary configurations, for example having three or more regular dietary elements.

Relative to dogs on a fresh food only diet, dogs fed dry plus canned food (OR=1.85, p<0.0001), dry food only (OR=1.45, p<0.0001), and dry plus fresh food (OR=1.3, p=0.03) were more likely to be overweight/obese. Dry plus canned food (OR=2.6, p<0.0001) and dry food only (OR=2.1, p<0.0001) were also risk factors for being obese, but not dry plus fresh food (p>0.05). Dogs fed raw food only were less likely to be overweight/obese (OR=0.5, p=0.005), but there was no effect on obesity alone (p>0.05).

#### Age

Given the strong relationship between body condition and age in both the univariate and multivariate models, we investigated this in further detail. We found that likelihood of overweightness peaked around age 8-10, decreasing with further aging (Figure 4, A). Likelihood of obesity alone showed a similar non-linear pattern. This relationship supports our use of higher-order polynomial age terms in the logistic regression models.

**Figure 4:**
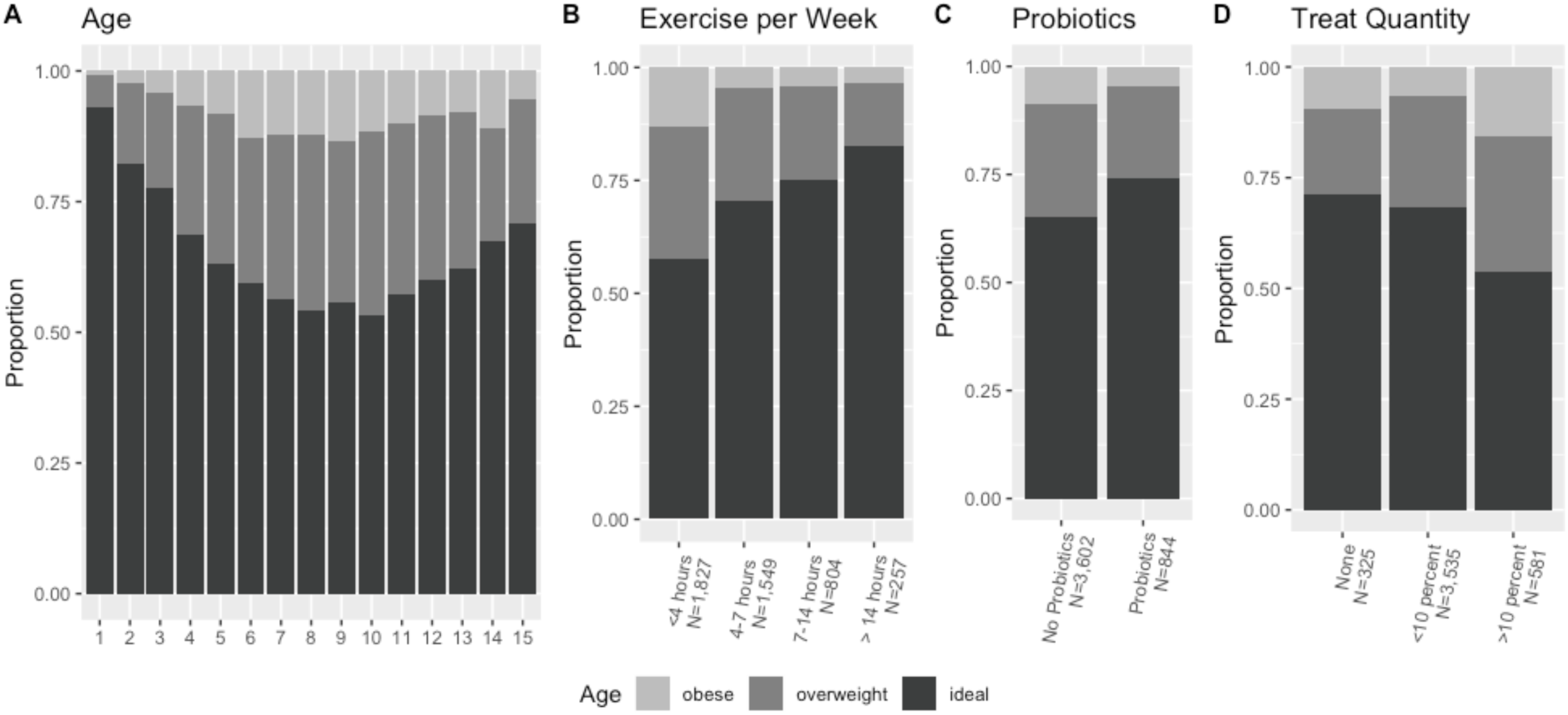
Factors Influencing Body Condition Scores. Proportion of overweight and obese dogs by age (A), exercise per week (B), probiotic consumption (C), and treat quantity (D).

#### Exercise

Since exercise was shown to be statistically significant in both the univariate and multivariate models (p<0.0001), we further considered this relationship (Figure 4, B). We found that incrementally increasing the amount of exercise per week decreased the likelihood of overweightness/obesity. The same pattern holds for obesity.

#### Probiotics

We also examined the relationship between probiotic supplementation and BCS (Figure 4, C). In total, 844 dogs were currently taking probiotics, and respondents reported using a broad range of commercial formulations. We found that probiotic supplementation was significant both with respect to overweight/obese status (OR=0.65, p<0.0001) and obese status (OR=0.46, p<0.0001) with dogs taking probiotics being more likely to be at an ideal weight. Given that dogs receiving probiotics may be more likely to be at a lower weight due to having a medical condition, we repeated this analysis using only the healthy cohort as previously defined (N=3,173). We found a significant relationship with both overweight/obese status (OR=0.65, p<0.0001) and obese status (OR=0.52, p = 0.002) in this cohort as well.

Since supplement usage may indicate increased owner health consciousness, we also performed a similar test with the other supplements included in our analysis in order to determine if this significant effect was specific to probiotics. We examined prebiotics (N=183), multivitamins (N=425), CBD oil (N=540), fish oil (N=812), herbal supplements (N=207), and immune support (N=121). None of these supplements showed a significant relationship with either overweight/obese status or obese status in chi-square tests (p>0.05).

#### Treats

Finally, we examined the univariate association between treat intake quantity (by percentage of caloric needs being met by treats) and BCS (Figure 4, D). Both the univariate and multivariate analyses confirmed that, while giving over 10% of a dog’s diet in treats was associated with higher BCS, there was no significant difference between giving under 10% of a dog’s diet in treats and abstaining completely, suggesting that giving treats in moderation is not a risk factor for either overweightness or obesity.

#### Non-Significant Factors

Given that some respondents (N=593) gave data for multiple dogs, and that joint households may exhibited shared environmental factors not captured by existent survey questions, we determined if there was a significant effect of household on reported body condition. We entered household into the logistic regression model as a random effect, and an analysis of deviance showed that it was not significant (p>0.05). Furthermore, given findings from other studies that identified a relationship between sex and BCS (e.g. (6), or an significant interaction between sex and neutered status on BCS (10), the analysis was repeated in the subset of animals for which sex data were provided (N=3,922) but sex was not significant either as a main effect or in interaction with neutering (p>0.05). Given previously reported overweightness rates of 52% and 41% for females and males respectively (5), and α=0.05, we had β>.99 power to detect a main effect, and following previously published guidelines for calculating power for interactions (16), we calculated that we had β>.90 power to detect an interaction between sex and neutering. Therefore, we are relatively confident in reporting that no relationship between sex and BCS exists in this sample.

## Discussion

This study employed a cross-sectional direct-to-consumer data collection approach to identify risk factors associated with increased owner-assessed body condition. Overweightness and obesity were common issues in this population, with 33% of dogs reported as overweight/obese and 8% reported as obese. This is comparable to the respective figures obtained in a study of 21,754 dogs in the United States wherein body condition scores were assessed by a veterinarian (3). The large dataset assembled here allowed us to identify and then validate risk factors using multiple approaches. Overweightness/obesity and obesity were each examined through three logistic regression models, with variables chosen through stepwise selection, Elastic Net selection, and an XGBoost machine learning model.

Many of the risk factors identified recapitulate robust findings from previous studies. Exercise has previously been identified as protective against obesity (11,12,17). Neutering has also been found to be a risk factor for overweightness but not obesity (6), and we find more significant relationships with the combined overweightness/obesity models than with the models for obesity alone. Though this result may be due to decreased power, this potential distinction in severity may be important for dog owners to recognize when weighing the potential outcomes of neutering. This relationship is corroborated by studies showing that neutering dogs results in lower daily energy requirements combined with higher food consumption (18). Sex has previously been found to be a risk factor both alone (6) and in interaction with neutering (10), though other studies did not identify a significant effect (9). We specifically tested for an association between sex and weight and, despite having adequate power, did not detect an effect in our sample. While a number of studies (4,5,8) have reported increased age as a risk factor for weight issues, we found in our sample that weight peaks around age 8-10 years, with older animals less likely to be overweight or obese than their middle-aged counterparts. This mirrors findings from other larger studies (3,5,6,9). This finding may reflect the decreased lifespans of overweight and obese dogs.

One novel finding of our study is the association between increased BCS and the consumption of dry food, which has previously been observed in cats (19, 20) but not in dogs (3, 11). Specifically, we found that, compared to fresh food, dry food is significantly associated with overweightness/obesity, both alone and in combination with canned or fresh food. Furthermore, dry food only diets and dry plus canned combination diets were significantly associated with obesity, but not dry plus fresh combination diets. This may indicate that supplementing with other types of food ameliorates the more obesogenic consequences of certain diets. The relationship between dry food and increased BCS may be partially due to the tendency of owners to inaccurately measure dry food portions (21), which may be alleviated by the decreased caloric density of fresh and raw foods. We posit that previous studies did not identify a relationship between dry food and increased body weight due to the lack of fresh and raw alternatives commercially available in past years. We also found that feeding commercial and home-prepared fresh, frozen, and raw foods is associated with a lower likelihood of being overweight or obese; however, diets containing raw animal products may contain higher rates of bacterial contamination than other foods and may pose health risks to companion animals (22). Observed differences in dietary factors could be due to a number of non-nutritional factors such as the provided portion sizes, the individualized portioning of some fresh diets, or the higher price point per calorie of some diets. Future studies should examine the effects of varied proportions of these dietary elements and experimental dietary manipulation to determine the direction of causality.

Another novel finding is that supplementation with probiotics is associated with being at an ideal weight. This finding is specific to probiotics among other supplements reported in the survey, and the effect persists in a sample composed of dogs without major reported health issues. This relationship is supported by work that shows significant differences in the gut microbiota of normal and obese dogs (23), as well as recent reviews and meta-analyses of experimental and clinical trials of probiotic supplementation in different species (24–26). However, to the best of our knowledge this is the first large cross-sectional demographic study that has specifically identified probiotic supplementation as a potential protective factor with regard to overweightness and obesity. The mechanisms behind this relationship are unclear, but in addition to modulating energy harvest and nutrient absorption through alterations in microbiota, probiotics might act through improving insulin sensitivity (27) or increasing satiety (24). Additional prospective data should be collected to identify whether there is a causal relationship.

We also identified treat feeding practices as robust predictors of body condition. In contrast to previous studies which identify even a moderate frequency of treat-feeding as a risk factor (8) we find that feeding treats in moderation, i.e. with 10% or less of total caloric needs being met by treats, is neither associated with overweightness/obesity nor obesity. This discordance between findings may be due to differential effects of treat quantity and treat frequency. However, these results should be interpreted cautiously as owner assessment of calories provided by a given treat may often be imprecise.

The presence of other dogs has been previously studied as a risk factor for increased BCS, with some studies identifying living in a single-dog household as a risk factor (11) and others finding no significant relationship (5). We find that the presence of other dogs in the household is associated with higher body weight, and we posit this may be due to the difficulty of implementing food restriction in the presence of a dog of healthy weight that is being concurrently fed.

Our models also included owner assessments of canine temperamental factors, including food motivation level, pet appetite, and overall mood. Significant relationships between obesity and temperamental factors such as appetite and food motivation have previously been identified (12,28,29). It is possible that dogs that express higher food motivation and appetite receive more calories from their owners, and food-seeking behavior may be associated with decreased owner compliance with weight loss regimens. While dogs’ subjective experience is difficult to measure, owner-perceived emotional eating in dogs has been associated with owner-assessed overall mood and anxiety (30). Tail-chasing, which emerged as a significant factor in some models, may also be an index of behavioral issues (31). While the links between behavior, mood, and obesity are unclear and likely to be multifactorial, serotonin has been implicated in both mood and satiety (32) and serotonin-reuptake inhibitors have been shown to decrease tail-chasing behavior (33).

Finally, we also note that our models identified additional variables, though it is difficult to pinpoint the statistical reasons for these differences. We identified increased dental treatment frequency as significant; however, the interpretation of this finding remains unclear. It is possible that increased dental visit frequency reflects higher health consciousness in owners or more frequent veterinary counseling on prevention of weight issues. A causal relationship between periodontal inflammation and obesity seems unlikely, but there could be effects on the microbiome or other processes that might contribute to adiposity. Some of our models also identified a significant association between rural environments and increased BCS, which is corroborated by previous studies (6, 9).

Limitations of this investigation include the use of owner assessment of BCS and owner report of other questionnaire items; prior studies have shown that owners inaccurately report body condition (34). However, we validated the use of BCS in this sample by cross-referencing BCS against owner-reported current-to-ideal body weight ratios and identified significant overlap as detailed in Methods. Another shortcoming is that we did not solicit demographic or lifestyle data of the owners, such as age and income, which have been previously shown to be associated with canine weight issues (8). Another potential limitation of our findings is selection bias, as our data were drawn primarily from the customer base of a pet health company providing individually portioned fresh meals, which may be more health-conscious than the aggregate dog owning population. However, this study is the first to report results from this unique population, broadly in accord with previously reported risk factors, supporting the generalizability of these findings. Finally, as the associations between the reported risk factors and BCS were cross-sectional, temporality and causality were not addressed. Future studies should seek to experimentally manipulate these risk factors individually or in combination to observe their potentially synergistic effects on weight loss or maintenance, particularly diet, probiotics, and treat quantity as they may be most easily manipulated by dog owners.

## Conclusions

In this cross-sectional, owner-reported study of 4,446 dogs, we identified risk factors for overweightness and obesity using stepwise selection, and then confirmed these findings using Elastic Net and XGBoost. Our findings recapitulated risk factors robustly identified by other studies, including exercise, neutering, and a non-linear contribution of age. We also further elucidated the contribution of previously studied risk factors; we identified dry food consumption, excessive treat intake, and canine temperament as risk factors. Finally, we identified probiotic supplementation as a potential protective factor. These findings are of clinical and practical significance because the demographic factors identified may be used to proactively identify dogs at increased risk for overweightness and obesity, and the lifestyle factors identified may guide the development of interventions. Further studies employing experimental manipulations are needed to establish whether these relationships are causal.

## Methods

### Data collection and selection procedures

Customers of NomNomNow, a pet food and health company, were asked to complete a comprehensive online health assessment for each pet in their care, composed of five broadly targeted questionnaires containing questions about signalment, overall wellness, diet and lifestyle, medical history, and product preferences. These included single-choice checkbox, multiple-choice radio-button, dropdown, and fill-in-the-blank questions. We collected survey responses from a total of 7,942 dogs of which 5,135 were complete. Dogs under one year of age (N=415), pregnant or lactating dogs (N=4), and underweight dogs (N=272) were excluded from the analysis, yielding a final sample of 4,446 dogs from 3,853 unique households broadly distributed across the United States. A maximum of 122 questions were asked regarding each dog, as some questions were served contingent upon responses to other questions. After dummy-coding and grouping, there were 372 individual features, which were reduced to 47 after removing variables related to medical conditions and treatment, product preferences, and highly correlated or redundant questions. This initial exclusion step was performed because Monte Carlo simulations show improved performance in stepwise models with fewer nuisance variables (35). After further removing variables that were more than 5% incomplete, we were left with 45 features which were used for subsequent analysis. The survey questions that contributed to the final features are detailed in [Supplementary File 6].

### Overweightness and Obesity

Weight status was assessed by dog owners using an image-based body condition score (BCS) chart accompanied by a verbal description. BCS has been shown to be strongly correlated with gold-standard dual-energy X-ray absorptiometry (DEXA) scans (36) and is regularly used both by veterinarians and researchers for clinical assessment of overweightness and obesity. Respondents were shown six options reflecting a nine point scale (BCS, Supplementary File 7 (13)). For the purposes of this analysis, we grouped the six pictures into four categories: underweight (corresponding to BCS 3 or lower); ideal (corresponding to BCS 4-5); overweight (corresponding to BCS 6); and obese (corresponding to BCS 7 or higher). Since owners have been reported to systematically misperceive body condition scores (34) we validated these scores against current and ideal body weights reported by a subset of respondents. Owner-reported BCS showed strong correspondence with owner-reported current-to-ideal body weight ratios. Using suggested ratio guidelines of >1.15 for overweightness and >1.30 for obesity (1) we calculated sensitivity (the proportion of overweight dogs by weight ratio correctly selected as overweight by BCS) and specificity (the proportion of ideal weight dogs by weight ratio correctly selected as ideal by BCS). We calculated 90% sensitivity and 79% specificity for overweightness/obesity, and 57% sensitivity and 94% specificity for obesity. Hence, dogs with BCS >= 6 are referred to as “overweight” and those with BCS >=7 as “obese.” For the survey image for body condition shown to respondents see Supplementary File 7.

### Diet

Owners were asked to check a box for each food type currently being fed, with multiple selections allowed. The final binned categories consisted of canned food, dry food (kibble), raw with home-prepared raw foods, dehydrated with freeze-dried foods, and commercial fresh/frozen with home-cooked foods. The seven most common combinations of diets were identified to be dry food only (N=938), canned food only (N=182), fresh food only (N=1,001), dry plus canned food (N=349), dry plus fresh food (N=376), dehydrated/freeze-dried only (N=109), raw only (N=136), and other configurations of these same elements (N=1,355); these diets were coded as factors and entered into the model as such. We did not ask for proportions of these dietary elements.

### Subject Characteristics

Out of a total of 4,446 dogs, 2,967 (67%) were at an ideal weight and 1,497 (33%) were overweight or obese, of which 1,142 (25% of total) were overweight and 355 (8% of total) were obese. In terms of sex, 1,727 (39%) were neutered male, 1,733 (39%) were neutered female, 262 (5.9%) were intact male, and 197 (4.4%) were intact female; 527 did not include both sex and neutered status. The average age was 6.74±4.09 years and the average weight was 13.6±11.8 kg.

### Data analysis

For each of the variable selection procedures, two models were run: the first to differentiate ideal weight dogs from overweight and obese dogs, and the second to differentiate ideal weight dogs from the smaller subset of dogs reported to be obese. The statistical significance for all variable selection procedures was set at α=0.05.

#### Univariate Analysis

Univariate associations between BCS and each of the 45 variables under consideration were examined. We performed chi-square tests for binary variables, t-tests for continuous variables, and ANOVA for categorical/ordinal variables, including arguably ordered factors such as exercise and treats, due to the possibility of non-linear effects. We did not correct for multiple comparisons in this analysis, in order to obtain a fuller, more descriptive, model of potential associations.

#### Method 1: Logistic Regression with Stepwise Selection

A logistic model was fit using variables selected through bidirectional stepwise selection using `StepAIC` in R. However, given the statistical issues raised around traditional stepwise selection, including the use of Monte Carlo simulations to show how infrequently all correct explanatory variables are selected during data mining (35), two additional methods were used to identify ensembles of risk factors, which were then entered into a traditional logistic regression to maximize interpretability.

#### Method 2: Elastic Net Selection

`ElasticNetCV` from the `sklearn` package was used to select the parameters λ and α, which control the level of penalty and the relative ratio of L_1_ and L_2_ penalties (39). Twenty iterations of Elastic Net were run using all variables, and the most important variables selected by Elastic Net were used to fit a logistic regression model to maximize interpretability. For both the overweight/obese and obese models, the best ensemble of risk factors was selected by consecutively adding variables and performing ANOVAs with significance tests to assess the contribution of each additional variable in the nested model, continuing the process until reaching a variable that did not significantly improve the model. Note that this differs from the true stepwise selection procedure performed before, in that the variables are added in order of importance in the complete model, rather than by their additive contribution to the partial model.

#### Method 3: XGBoost Selection

A grid search using `GridSearchCV()` was used to identify parameter values. Twenty iterations of XGBoost were run using all variables, and variable importance was averaged across iterations. The most important variables were entered stepwise into a logistic regression model until subsequent entered variables failed to significantly improve the predictive ability of the model as detailed for Elastic Net.

#### Data Imputation

Since not all survey questions were required, the 45 variables used in the analysis had between 0-4.5% missing data, which were inferred via imputation using `MissForest()` in R (40). Missingness and out-of-box imputation accuracy are provided for each of the variables in [Supplementary File 8]. Variables with over 5% missingness were excluded from analysis. Once variables were identified as important through the variable selection methods delineated above, the final logistic regression model was fit using only complete cases so as to avoid inaccurate or biased model fit values.

Univariate analysis, imputation, stepwise selection, and logistic regressions were conducted in R version 3.5.2. Elastic Net and XGBoost were conducted in Python version 3.6.7.

## Declarations

### Ethics statement

This study was carried out in accordance with the recommendations of the Federal Policy for the Protection of Human Subjects, U.S. Department of Health and Human Services. Since our study gathered information from owners focused solely on their dogs it does not meet the definition of human subjects research and therefore did not require review by an Institutional Review Board.

### Consent for publication

Survey respondents agreed to a Terms of Service (TOS) consenting to using pet information for research and publication. Furthermore, all the data in this study has been deidentified.

### Availability of data and materials

The deidentified datasets generated and/or analyzed during the current study are not publicly available since they are generated for proprietary business and marketing purposes, but are available from the corresponding author upon reasonable request.

### Authors’ contributions

LP conceived the scope of the work, analyzed and interpreted the data, drafted the manuscript, and created figures and tables. JS designed the survey questions and reviewed the findings and manuscript. JT interpreted the data and critically reviewed the manuscript. DM implemented the owner-reported assessment questions. RH conceived the scope of the work, designed the survey questions, and reviewed the findings and the manuscript. AJ conceived the scope of the work, interpreted the data, and reviewed the findings and the manuscript.

## Acknowledgements

The authors would like to thank the NomNomNow, Inc. Scientific Advisory Board for their thoughtful consideration and helpful discussion regarding data analysis and interpretation. We are grateful to our survey participants for providing us with the data on their dogs.

