## Supplementary figures and images for "Risk factors associated with canine overweightness and obesity in an owner-reported survey"

### Supplementary File 1

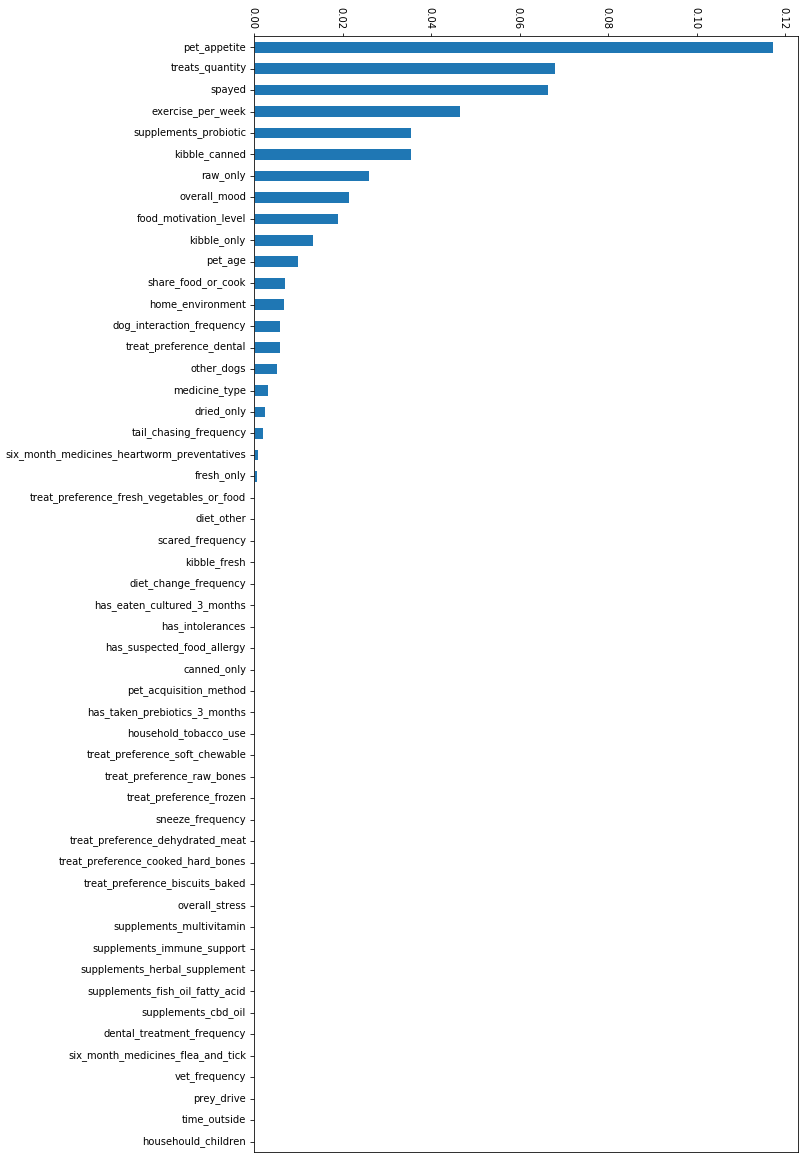

### Supplementary File 2

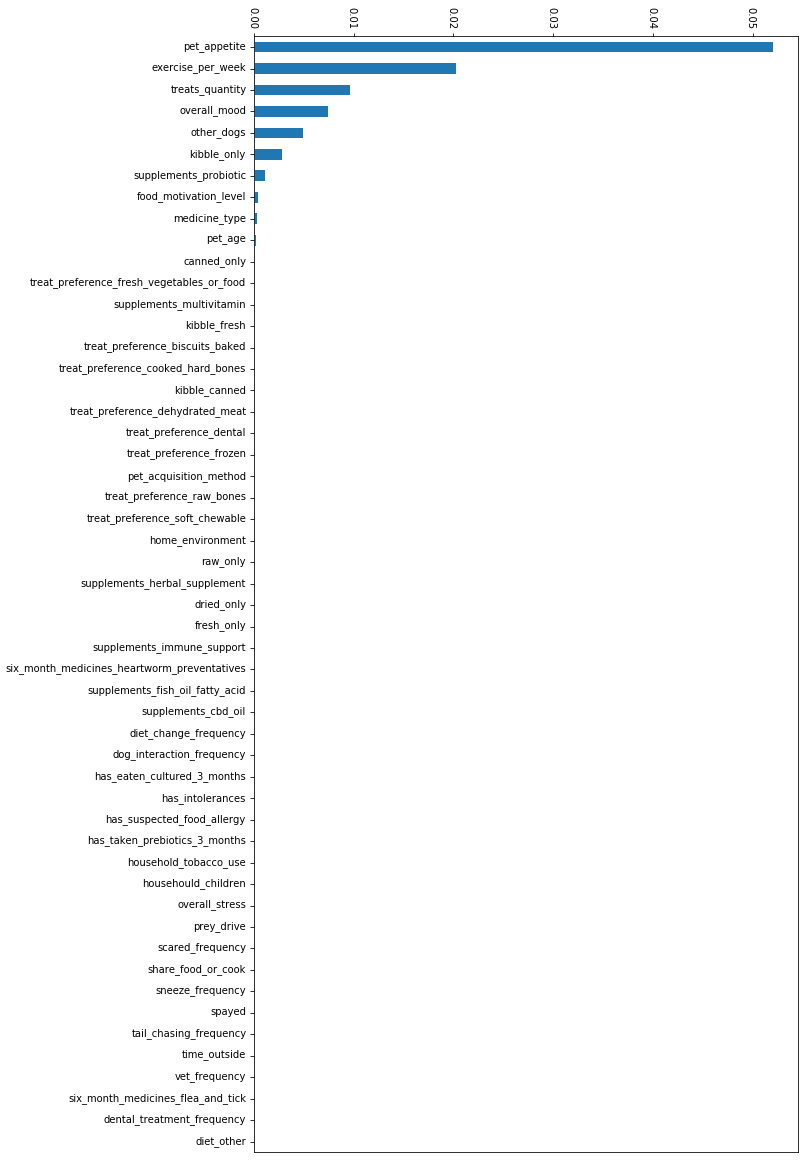

### Supplementary File 3

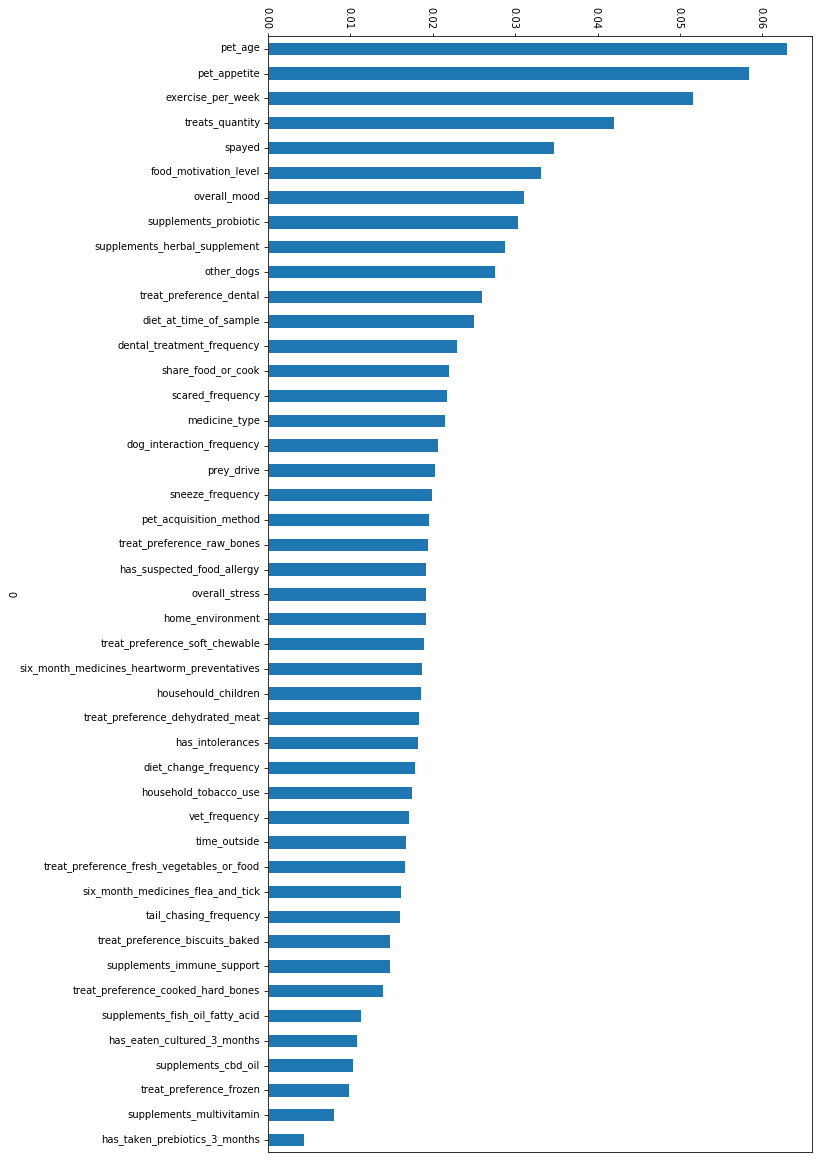

### Supplementary File 4

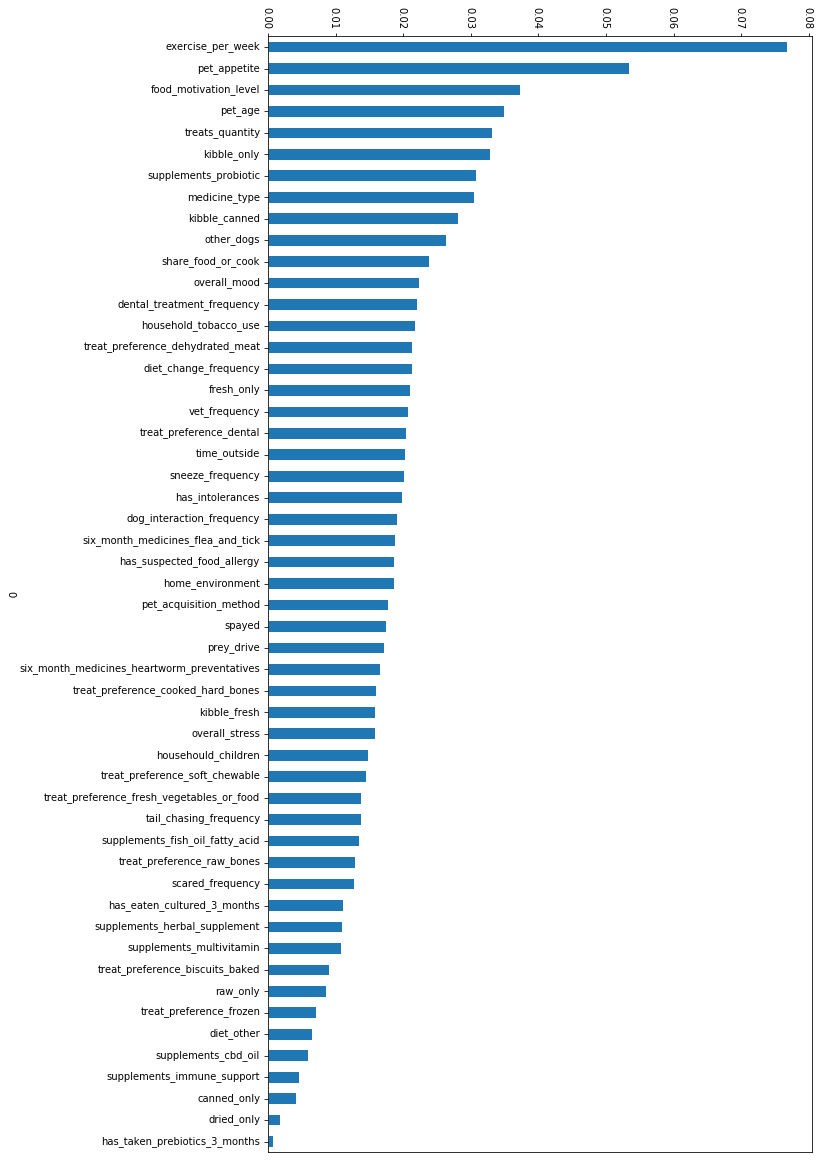

### Supplementary File 7

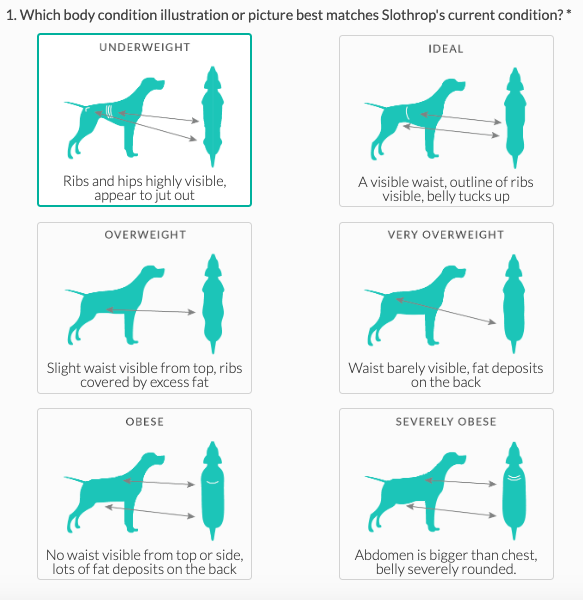
