## Supplementary File 5 for "Risk factors associated with canine overweightness and obesity in an owner-reported survey"

| Table 4: Final Multivariable Logistic Regression Models for the Subgroup Without Major Medical Conditions | | | | | |
| --- | --- | --- | --- | --- | --- |
|  |  | Overweight and Obese (N=1004) vs Ideal (N=1977) | | Obese (N=246) vs Ideal (N=1977) | |
| Feature |  | OR [95% CI] | p-value | OR [95% CI] | p-value |
| Age | Years | 1.64 [1.50-1.80] | <0.0001*** | 1.95 [1.61-2.37] | <0.0001*** |
|  | Years^2^ [A] | 0.97 [0.97-0.98] | <0.0001*** | 0.96 [0.95-0.98] | <0.0001*** |
| Exercise per week | <4 hours | *Reference* | | *Reference* | |
|  | 4-7 hours | 0.59 [0.48-0.71] | <0.0001*** | 0.27 [0.18-0.39] | <0.0001*** |
|  | 7-14 hours | 0.51 [0.40-0.65] | <0.0001*** | 0.24 [0.14-0.40] | <0.0001*** |
|  | >14 hours | 0.38 [024-0.59] | <0.0001*** | 0.19 [0.07-0.50] | 0.0007*** |
| Diet Composition | Fresh | *Reference* | | *Reference* | |
|  | Dry | 1.48 [1.16-1.88] | 0.0001*** | 2.16 [1.39-3.36] | 0.0006*** |
|  | Canned | 0.98 [0.63-1.54] | >0.05 | 1.30 [0.58-2.90] | >0.05 |
|  | Dehydrated/ Freeze-Dried | 0.72 [0.41-1.25] | >0.05 | 0.48 [0.14-1.65] | >0.05 |
|  | Raw | 0.61 [0.36-1.04] | >0.05 | 0.44 [0.14-1.43] | >0.05 |
|  | Kibble and Canned | 2.19 [1.59-3.03] | 0.0001*** | 3.08 [1.75-5.42] | <0.0001*** |
|  | Kibble and Fresh | 1.48 [1.08-2.03] | 0.02* | 1.54 [0.81-2.93] | >0.05 |
|  | Other Combination | 1.01 [0.79-1.31] | >0.05 | 0.99 [0.58-1.67] | >0.05 |
| Food Motivation Level | 1-5 scale | 1.19 [1.10-1.28] | <0.0001*** | 1.31 [1.13-1.52] | 0.0003*** |
| Other Dogs |  | ns | ns | 1.86 [1.34-2.59] | 0.0002*** |
| Overall Mood | 1-5 scale | 1.26 [1.15-1.39] | <0.0001*** | 1.41 [1.19-1.67] | <0.0001*** |
| Pet Appetite | 1-5 scale | 2.13 [1.73-2.62] | <0.0001*** | 4.83 [3.19-7.29] | 0.0001*** |
| Share Food or Cook |  | 1.24 [1.03-1.48] | 0.02* | ns | ns |
| Neutered |  | 2.12 [1.45-3.09] | <0.0001*** | 3.30 [1.45-7.52] | 0.004** |
| Dental Treatment Frequency | 1-3 scale | 0.77 [0.68-0.88] | <0.0001*** | 0.68 [0.54-0.87] | 0.002** |
| Tail Chasing Frequency | 1-5 scale | ns | ns | ns | ns |
| Treats Quantity | None | *Reference* | | *Reference* | |
|  | Less than 10% | 1.15 [0.81-1.64] | >0.05 | 0.98 [0.53-1.83] | >0.05 |
|  | More than 10% | 2.50 [1.66-3.76] | <0.0001*** | 5.36 [2.66-10.79] | <0.0001*** |
| Supplements Probiotic |  | 0.72 [0.57-0.92 | 0.01* | 0.60 [0.37-0.99] | 0.04* |
| Medicine Type | Conventional | ns | ns | ns | ns |
|  | Integrative | ns | ns | ns | ns |
|  | Holistic | ns | ns | ns | ns |
| Home Environment | Urban | *Reference* | | *Reference* | |
|  | Suburban | 1.17 [0.95-1.44] | >0.05 | ns | ns |
|  | Rural | 1.35 [1.02-1.79] | 0.03* | ns | ns |

OR: Odds Ratio; 95% CI: 95% Confidence Interval; Food Motivation Level: 1-Not at all, 5-Very; Mood: 1-Excellent, 5-Depressed; Pet Appetite: 1-Poor, 5-Excellent; Dental Frequency: 1-Never, 3-Three or more times; Tail Chasing Frequency: 1-Never, 5-Multiple times per day

[A] Age as a quadratic term was eliminated from the confirmatory variable selection procedures to maximize interpretability. Note also that XGBoost already handles nonlinearities in the input features through the potential selection of multiple decision points per variable.

[B] The coefficient of quadratic terms denotes the curvature. In this case the negative coefficient indicates that the curve is concave, with dogs in the middle of the age range at highest risk.
