## Supplementary File 6 for "Risk factors associated with canine overweightness and obesity in an owner-reported survey"

| **Question** | **Question Name** | **Type of questions** | **Mandatory** | **Choices** |
| --- | --- | --- | --- | --- |
| Birthday day | birthday_day | Single selection | No |  |
| Birthday month | birthday_month | Single selection | No |  |
| Birthday year | birthday_year | Single selection | No |  |
| Gender | gender | Single selection | No | Male; Female. |
| Which body condition illustration or picture best matches Slothrop’s current condition? | body_condition | Single selection | Yes | Ribs and hips highly visible, appear to jut out;  A visible waist, outline of ribs visible, belly tucks up;  Slight waist visible from top, ribs covered by excess fat;  Waist barely visible, fat deposits on the back;  No waist visible from top or side, lots of fat deposits on the back;  Abdomen is bigger than chest, belly severely rounded. |
| Does Slothrop have any of the following medical issues (select all that apply)? | medical_issues | Multiple selection | Yes | Allergies;  Arthritis, Joint Pain or abnormalities;  Behavioral issues (e.g. anxiety, aggression);  Cancer;  Chronic diarrhea;  Chronic ear infections;  Chronic eye issues (e.g. watery/dry eyes, glaucoma);  Chronic vomiting;  Cushing’s disease (hyperadrenocorticism);  Diabetes;  Disc, neck, or back problems;  Heart disease or failure;  Kidney disease or failure;  Liver disease;  Obesity or overweight;  Oral issues: Bad teeth, bad breath, dental disease;  Pancreatitis (now or previously);  Seizures or epilepsy;  Thyroid disorder (e.g. hyperthyroid, hypothyroid);  Urinary infections or inflammation (e.g. cystitis);  Urinary stones (now or previously);  None of the above. |
| Is Slothrop currently pregnant or lactating? | currently_pregnant_or_lactating | Single selection | Yes | Yes; No. |
| How would you describe Slothrop's diet in the two weeks prior to collecting their sample? | diet_at_time_of_sample | Multiple selection | Yes | Kibble; Raw; Canned; Home-prepared raw; Home-prepared cooked; Commercial fresh; Commercial frozen; Freeze dried; Dehydrated; Other. |
| Treats provide about how much of Slothrop’s overall food intake? | treats_quantity | Single selection | Yes | None; 10% or less; >10%. |
| How often do you change Slothrop's diet? | diet_change_frequency | Single selection | Yes | Daily;  About weekly;  About monthly;  A couple of times a year;  Rarely. |
| Did you share your food or cook for Slothrop even if not the whole of the diet before feeding NomNomNow? | share_food_or_cook | Single selection | Yes | Yes; No. |
| Do you give Slothrop any of the following supplements | supplements | Multiple selection | Yes | Allergies; Behavior; CBD oil; Calming/Anxiety; Cognitive Development; Digestion; Eye health; Fish oil; Heart health; Herbal; Immune support; Joint & mobility; Multivitamin; Other; Probiotics; Skin & coat; Wegiht management; None of the above. |
| How often do you normally take Slothrop to the veterinarian? | vet_frequency | Single selection | No | Only if there is a problem; Once yearly; Twice yearly; More than twice yearly. |
| Which of the following best describes your approach to medical care for Slothrop's health? | medicine_type | Single selection | Yes | Conventional only (phamaceutical drugs, conventional veterinarian); Integrative (mixture of conventional and alternative/holistic therapies); Holistic only (natural supplements, diets, general avoidance of drugs and vaccines). |
| Has Slothrop been spayed or neutered? | spayed | Single selection | Yes | Yes; No. |
| Has Slothrop taken any of the following medications within the last 6 months (check all that apply) | six_month_medicines | Multiple selection | Yes | Antibiotics (e.g. Clavamox, cephalexin, Baytril); Antifungal (ketoconazole, itraconazole); Antiparasitics (pyrantel, Panacur, praziquantal (not routine monthly preventatives)); Behavior modifying drugs and sedatives (Clomicalm, Prozac, trazadone, acepromazine); Chemotherapeutics (“chemo”); Flea and tick medications (topical or oral); Heartworm preventatives; Medications for vomiting or diarrhea (metronidazole, Cerenia); Non-steroidal anti-inflammatories (NSAIDS eg. Rimadyl, Dermaxx, Previcox, Metacam or generics); Opioid or opioid-like pain relievers (hydrocodone, tramadol); Other medications; Steroids (prednisone, prednisolone); Thyroid supplement; None of the above. |
| Which antibiotics has Slothrop taken in the last 6 months? | six_month_antibiotics | Fill in the blank | No |  |
| Please list the other medicines has Slothrop been prescribed by their vet in the last 6 months? | six_month_other_medicines | Fill in the blank | No |  |
| Does your household have children? | househould_children | Single selection | Yes | Yes; No. |
| How frequently does Slothrop interact with other dogs (outside of your household)? | dog_interaction_frequency | Single selection | Yes | Routinely (dog attends daycare, playgroup, group walks etc. on a regular basis, ≥1/week); Occasionally (dog has playdates, or visits dog park occasionally, ≥1/month); Rarely (interacts with other pets in passing); Never. |
| Do you have other pets or animals? | other_pets | Multiple selection | Yes | Dogs; Cats; Pocket pets (e.g. hamster, gerbil, etc.); Birds; Farm animals; No other pets or animals. |
| How many other dogs do you have? | other_dogs | Single selection | No |  |
| How would you describe your pet's home environment? | home_environment | Single selection | Yes | Urban; Suburban; Rural. |
| How much time does Slothrop spend outside? | time_outside | Single selection | Yes | Never outside; 0-1 hours/day; 1-4 hours/day; 4-8 hours/day; >8 hours/day. |
| Does anyone in the household smoke tobacco? | household_tobacco_use | Single selection | Yes | Yes; No. |
| Through which method did Slothrop join your family? | pet_acquisition_method | Single selection | Yes | Rescue or adoption; Breeder; Found them; Gift from friend or family; Other purchase; Other. |
| How much exercise does Slothrop get each week? | exercise_per_week | Single selection | Yes | <4 hours; 4-7 hours; 7-14 hours; 14 or more hours;, |
| Does Slothrop have any food intolerances? | has_intolerances | Single selection | Yes | Yes; No. |
| Do you suspect Slothrop has a food allergy? | has_suspected_food_allergy | Single selection | Yes | Yes; No. |
| What kind of treats does Slothrop prefer? | treat_preference | Multiple selection | No | Biscuits or baked treats; Fresh vegetables or foods; Soft chewable treats; Dental treats or chews; Raw bones; Cooked or hard bones; Dehydrated meat treats; Frozen treats; None of the above. |
| Does Slothrop eat poop? | is_poop_eater | Single selection | Yes | Yes; No; |
| Is Slothrop taking, or has taken, probiotics within the last three months (products containing live cultures)? | has_taken_probiotics_3_months | Single selection | Yes | Yes; No. |
| Has Slothrop consumed any cultured or fermented food (yogurt, kimchi, etc.) within the last three months? | has_eaten_cultured_3_months | Single selection | Yes | Yes; No. |
| Is Slothrop taking, or has taken, prebiotics (such as soluble fiber, FOS, inulin/chicory root) within the last three months? | has_taken_prebiotics_3_months | Single selection | Yes | Yes; No. |
| How nervous or scared is Slothrop around new people? | scared_frequency_new_people | Single selection | Yes | Very; Somewhat; Average; Mildly; Not at all. |
| How nervous or scared is Slothrop in new surroundings? | scared_frequency_new_surroundings | Single selection | Yes | Very; Somewhat; Average; Mildly; Not at all. |
| How nervous or scared is Slothrop around unfamiliar pets? | scared_frequency_unfamiliar_pets | Single selection | Yes | Very; Somewhat; Average; Mildly; Not at all. |
| How frequently does Slothrop chase other animals (i.e. How much prey drive)? | prey_drive | Single selection | Yes | Never; Rarely; Occasionally; Often; Very often. |
| How frequently does Slothrop mount inappropriately? | inappropriate_mounting_frequency | Single selection | Yes | Never; Rarely; Occasionally; Often; Very often. |
| How would you rate Slothrop’s overall aggression? | overall_aggression | Single selection | Yes | Very aggressive; Somewhat aggressive; Average aggressive level; Mildly aggressive; Not aggressive. |
| How aggressive is Slothrop towards other pets? | other_pet_aggression | Single selection | Yes | Very; Somewhat; Average; Mildly; Not at all. |
| How aggressive is Slothrop towards unfamiliar people? | unfamiliar_people_aggression | Single selection | Yes | Very; Somewhat; Average; Mildly; Not at all. |
| How would you rate Slothrop’s overall mood? | overall_mood | Single selection | Yes | Excellent; Very good; Good; Fair; Depressed. |
| How frequently does Slothrop sneeze? | sneeze_frequency | Single selection | Yes | Never; Multiple times each day; Daily; Weekly; Monthly. |
| How frequently does Slothrop chase their tail? | tail_chasing_frequency | Single selection | Yes | Never; Multiple times each day; Daily; Weekly; Monthly. |
| How frequently does Slothrop mark/territorially pee? | mark_territory_frequency | Single selection | Yes | Never; Multiple times each day; Daily; Weekly; Monthly. |
| How would you describe Slothrop's appetite? | pet_appetite | Single selection | Yes | Increased; Average; Decreased |
