## Supplementary File 8 for "Risk factors associated with canine overweightness and obesity in an owner-reported survey"

| **factor_name** | **missingness** | **error_rate** | **error_type** |
| --- | --- | --- | --- |
| pet_age | 0 | NaN | MSE |
| dental_treatment_frequency | 93 | 0.411469224 | MSE |
| diet_change_frequency | 7 | 0.86676793 | MSE |
| dog_interaction_frequency | 15 | 0.723309284 | MSE |
| exercise_per_week | 9 | 0.584506573 | MSE |
| food_motivation_level | 7 | 1.418402928 | MSE |
| has_eaten_cultured_3_months | 16 | 0.188261851 | PFC |
| has_intolerances | 26 | 0.152488688 | PFC |
| has_suspected_food_allergy | 13 | 0.154974058 | PFC |
| has_taken_prebiotics_3_months | 13 | 0.041055718 | PFC |
| household_tobacco_use | 21 | 0.073446328 | PFC |
| househould_children | 21 | 0.161355932 | PFC |
| other_dogs | 0 | 0 | PFC |
| overall_mood | 10 | 0.69416929 | MSE |
| overall_stress | 5 | 1.170181181 | MSE |
| pet_appetite | 202 | 0.195735851 | MSE |
| prey_drive | 13 | 1.549221874 | MSE |
| scared_frequency | 24 | 1.207320567 | MSE |
| share_food_or_cook | 10 | 0.350090171 | PFC |
| sneeze_frequency | 26 | 1.545294547 | MSE |
| spayed | 18 | 0.095754291 | PFC |
| tail_chasing_frequency | 12 | 0.517255589 | MSE |
| time_outside | 113 | 0.303222353 | MSE |
| treats_quantity | 5 | 0.18813262 | MSE |
| vet_frequency | 0 | NaN | PFC |
| six_month_medicines_flea_and_tick | 0 | NaN | PFC |
| six_month_medicines_heartworm_preventatives | 0 | NaN | PFC |
| supplements_cbd_oil | 0 | NaN | PFC |
| supplements_fish_oil_fatty_acid | 0 | NaN | PFC |
| supplements_herbal_supplement | 0 | NaN | PFC |
| supplements_immune_support | 0 | NaN | PFC |
| supplements_multivitamin | 0 | NaN | PFC |
| supplements_probiotic | 0 | NaN | PFC |
| treat_preference_biscuits_baked | 0 | NaN | PFC |
| treat_preference_cooked_hard_bones | 0 | NaN | PFC |
| treat_preference_dehydrated_meat | 0 | NaN | PFC |
| treat_preference_dental | 0 | NaN | PFC |
| treat_preference_fresh_vegetables_or_food | 0 | NaN | PFC |
| treat_preference_frozen | 0 | NaN | PFC |
| treat_preference_raw_bones | 0 | NaN | PFC |
| treat_preference_soft_chewable | 0 | NaN | PFC |
| home_environment | 14 | 0.408619134 | PFC |
| medicine_type | 47 | 0.389406683 | PFC |
| pet_acquisition_method | 0 | NaN | PFC |
| diet_at_time_of_sample | 0 | NaN | PFC |
| MSE: mean squared error |  |  |  |
| PFC: proportion falsely classified |  |  |  |
| NaN: not applicable |  |  |  |
